# New insights into nCOVID-19 binding domain and its cellular receptors

**DOI:** 10.1101/2020.09.06.285023

**Authors:** Ankush Garg, Gaurav Kumar, Sharmistha Sinha

**Author notes:** Equal contribution (Ankush Garg, Gaurav Kumar).

## Abstract

nCOVID-19 virus makes cellular entry using its spike protein protruding out on its surface. Angiotensin converting enzyme 2 receptor has been identified as a receptor that mediates the viral entry by binding with the receptor binding motif of spike protein. In the present study, we elucidate the significance of N-terminal domain of spike protein in spike-receptor interactions. Recent clinical reports indicate a link between nCOVID-19 infections with patient comorbidities. The underlying reason behind this relationship is not clear. Using molecular docking, we study the affinity of the nCOVID-19 spike protein with cell receptors overexpressed under disease conditions. Our results suggest that certain cell receptors such as DC/L-SIGN, DPP4, IL22R and ephrin receptors could act as potential receptors for the spike protein. The receptor binding domain of nCOVID-19 is more flexible than that of SARS-COV and has a high propensity to undergo phase separation. Higher flexibility of nCOVID-19 receptor binding domain might enable it to bind multiple receptor partners. Further experimental work on the association of these receptors with spike protein may help us to explain the severity of nCOVID-19 infection in patients with comorbidities.

## Introduction

Early in the year 2020, a new viral disease began generating headlines throughout the world. The infection which started in the city of Wuhan, China soon spread across continents affecting more than 14 million people in 216 countries(https://www.who.int/emergencies/diseases/novel-coronavirus-2019/situation-reports/). Its symptoms are similar to those of common flu, such as cough, high temperature and shortening of breath in severe cases. By march 2020, World health organization declared it to be a pandemic. The decryption of the viral genome and phylogenetic analysis showed that the new virus (nCOVID-19) was similar to the bat derived SARS like coronaviruses such as SARS-COV with few genotypic differences (Andersen, Rambaut, Lipkin, Holmes, & Garry, 2020).

Recent clinical reports on COVID-19 show a link between patient comorbidity and nCOVID-19 infection ((Dai et al., 2020; Fang, Karakiulakis, & Roth, 2020; Hussain, Bhowmik, & do Vale Moreira, 2020; Liang et al., 2020; Passaro et al., 2020; Remuzzi & Remuzzi, 2020; Yang et al., 2020; Zhou et al., 2020). It has been found that patients with other comorbidities like diabetes and cancer are at higher risk during the COVID-19 pandemic. It is therefore reasonable to ask what makes certain individuals with comorbidities vulnerable to nCOVID-19 infection. To answer this question, we need a thorough understanding regarding the mechanism of viral entry into the cells. Structural characterization of the nCOVID-19 suggests, that similar to SARS-COV, the novel virus makes entry into human cells by interacting with the angiotensin converting enzyme (ACE2) receptors on cell surface. The interaction occurs via spike protein projecting out on the viral surface. The crystal structure of the nCOVID-19 spike protein has been solved in two independent studies (Walls et al., 2020; Wrapp et al., 2020). This spike protein exists as homo-trimer. Each monomer consists of domains S1 and S2 (**Fig 1a**). The receptor binding domain (RBD), residues C336-L518 present in the S1 domain of the spike protein shows strong affinity towards ACE2 and is suggested to play a crucial role in viral entry into the cells. The receptor binding motif (RBM), residues S438-Q506 in the RBD region (**Fig 1a**) interacts directly with the ACE2 during spike protein-ACE2 interaction. A useful strategy to prevent viral entry into the cells would be to target this RBM region of the spike protein with a potential inhibitor (Y. Han & Král, 2020).

**Fig 1:**
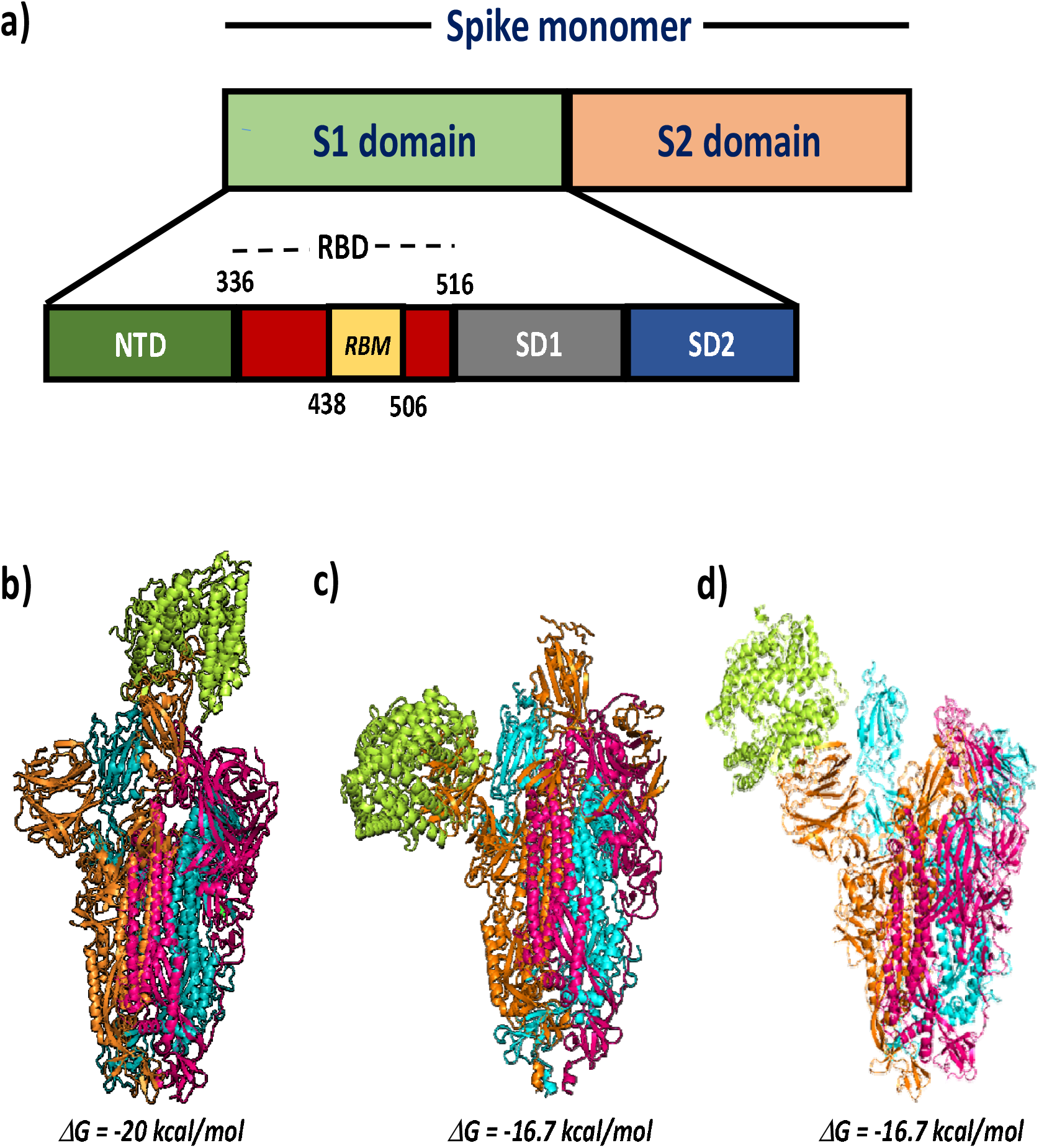
a) Schematic representation of nCOVID-19 spike protein monomer; ClusPro models showing interaction of ACE2 receptor (PDB ID: 1R42, in Limon) with b) SARS-COV spike protein (PDB ID: 6CRZ), c) COVID-19 spike protein (PDB ID: 6VSB) and d) SARS-COV spike protein-RBM deleted; Free energy of binding is calculated using PRODIGY server.

However, previous studies on Coronaviruses have shown that receptors other than ACE2 could also mediate viral entry into the cells. For example, reports on SARS-COV and MERS reveal the role of cell receptors such as DC/L-SIGN and DPP4 in viral cell entry (D. P. Han, Lohani, & Cho, 2007; Jeffers et al., 2004; Li et al., 2020; Marzi et al., 2004; Song et al., 2014). Interestingly, the SARS-COV spike protein residues involved in binding with DC/L-SIGN receptors are distinct from those present in the RBD region of the spike protein (D. P. Han et al., 2007). This raises a question if ACE2 is the only cell receptor that contributes to nCOVID-19 cell entry and whether targeting only the RBM region of the spike protein will be sufficient to stop the viral entry in to the cells. In this work, we address these questions using *in silico* studies. We first highlight the importance of spike residues beyond the RBM region in mediating spike receptors interactions. We also select few cell receptors based on their reported link with previously identified coronaviruses and their dysregulated expressions in cancer and diabetic patients. Using molecular docking, we look at the interactions of the n-COVID-19 spike protein trimer (PDB ID:6VSB) with the selected cell receptors. We also compare their affinities towards the RBD region of the spike protein. Our docking results help us to identify the potential cell receptors for nCOVID-19 spike protein.

## Material and methods

### Retrieval of protein crystal structure

The crystal structure of spike protein of the nCOV-19 and all cellular receptors are retrieved from RSCB database. The PDB ID of spike protein used for the protein-protein docking studies is 6VSB (nCOV-19) and 6CRZ (SARS-COV). The PDB ID of various receptors are 1R42 (ACE2), 4L72 (chain A corresponds to DPP4), 3DGC (chain R corresponds to IL22-R), 6GHV (DC-SIGN), 1XAR (L-SIGN), 3C8X (EphA2) and 3ETP (EphB2). We have removed all the water molecules before using them for docking studies.

### Protein-protein Docking

The docking between Spike protein/RBD with cell receptors is performed using ClusPro docking server (Kozakov et al., 2017). The docked model with the maximum number of clusters is selected for further analysis.

### Analysis of the Docked models

The binding affinity of the docked complexes were determined using PRODIGY server (Vangone & Bonvin, 2015; L. C. Xue, Rodrigues, Kastritis, Bonvin, & Vangone, 2016). The residues involved in interaction were identified using PDBSUM server (Laskowski, Jabłońska, Pravda, Vařeková, & Thornton, 2018). The H-bonds and salt bridges within the distance of 3Å were selected. We further predicted the total binding energy and atomic contact energy using FIREDOCK protein-protein docking refinement server(Andrusier, Nussinov, & Wolfson, 2007; Mashiach, Schneidman-Duhovny, Andrusier, Nussinov, & Wolfson, 2008).

### Prediction of disorderedness and phase separation

We predicted the structural disorderedness of spike protein of nCOVID-19 and SARS-COV using PONDR server(B. Xue, Dunbrack, Williams, Dunker, & Uversky, 2010). The phase separation propensity was predicted using CatGRANULE server(Bolognesi et al., 2016).

## Results

### In the absence of RBM residues, ACE2 binds to N-terminal domain of spike protein

To have a clear understanding regarding the residue level interactions between the spike protein and different cell receptors, we thought it would be better if we use the biological assembly of the spike protein (trimer) in our *in-silico* studies. Careful analysis of the COVID-19 spike protein structure (PDB ID: 6VSB) reveals that a significant portion of the RBM region is not solved. We wonder if the absence of the RBM residues impairs its association with the receptor ACE2. To check this, we perform protein-protein docking between ACE2 and nCOVID-19 spike protein trimer, using ClusPro docking server (Kozakov et al., 2017). For comparison with the SARS-COV, we also perform docking simulation between ACE2 and SARS-COV spike protein trimer (PDB ID: 6CRZ). In case of SARS-COV, the receptor ACE2 binds to the RBD region of spike protein with a free energy (ΔG) value of −20 kcal/mol. (**Fig 1b**). Since the RBD region in SARS-COV spike protein is completely modeled, the RBM residues are predicted to participate in both hydrogen bonding (H-bond) and salt bridges with ACE2 **(Supporting file, Table S1a-b)**. The receptor ACE2 interacts with chain A of the spike protein via 19 H-bonds and 6 salt bridges.

In case of nCOVID-19 spike protein, the receptor ACE2 binds to the spike protein with a ΔG value of −16.7 kcal/mol **(Fig 1c)**. The absence of the RBM residues in the solved crystal structure of nCOVID-19 spike protein does not impair its association with the ACE2 receptor. The receptor forms 17 H-bonds and 4 salt bridges with the N-terminal domain (NTD) of the chain A of spike protein trimer **(Supporting file, Table S2a-b**). This suggests that although ACE2 specifically binds to the RBM region of the spike protein, the absence of the RBM residues in spike protein may only lower its affinity towards ACE2 and not disrupt it. To Validate our observation, we delete the RBM residues (440-495) from the RBD region of SARS-COV spike protein (PDB ID: 6CRZ) and check its interaction with the ACE2. As shown in **Fig 1d**, the receptor binds with the NTD of the spike protein (ΔG = −16.7 kcal/mol). While the chain B of spike protein forms 1 H-bond with the receptor, the NTD of chain C forms 16 H-bonds and 3 salt bridges with the receptor ACE2 **(Supporting file, Table S3a-c)**. This suggests that the affinity of the ACE2 receptor towards residues beyond the RBM region must be taken into consideration while developing targeted therapeutics against the spike protein.

### Predicting receptors for nCOVID-19 spike protein other than ACE2

Post SARS-COV outbreak in 2002, many reports suggested the role of receptors DC-SIGN/L-SIGN in mediating viral entry into the cells (Jeffers et al., 2004; Marzi et al., 2004). They are C-type lectins with extracellular carbohydrate recognition domain (CRD) used for binding carbohydrate molecules. While one report showed that these receptors only augment ACE2 mediated viral infection, another study confirmed that DC/L-SIGN allow viral entry into the cells independently of the ACE2. To understand the affinity of these receptors towards the nCOVID-19 spike protein, we perform docking simulation between the spike protein and the carbohydrate recognition domain (CRD) of DC-SIGN (PDB ID: 6GHV)/L-SIGN (PDB ID: 1XAR) receptors. The CRD of both DC-SIGN and L-SIGN interact with the spike protein with a ΔG value of −14.3 and −14.2 kcal/mol respectively. (**Fig 2a & 2b**). In case of DC-SIGN, the residues Thr-333, Asn-334, Asn-354, Arg-357, Asn-360, chain C of spike protein participate in H-bonding with the receptor. **(Supporting file, Table S4a)**. While in case of L-SIGN, three residues from chain A and five residues from NTD of chain B form H-bonds with the receptor. Besides, Asp-364 and Arg577 from chain A and Glu-169 and Asp-228 from chain B are involved in salt bridge formation with receptor. (**Supporting file, Table S5a-d**). As the RBM residues are missing in the Spike protein structure used in the study, we perform docking interactions between the RBD region of the spike protein and the receptors. We compare the affinities of DC/L-SIGN receptors towards RBD region with that of ACE2. The ACE2 binds to the RBD with a ΔG value of −11.9 kcal/mol. The receptors D-SIGN and L-SIGN show similar affinity towards the RBD region with ΔG values of −10.7 and −9.8 kcal/mol respectively **(Supporting file, Fig S1a and S1b)**. It must be noted that the RBD residues involved in DC/L-SIGN binding are distinct from those involved in ACE2 binding **(Table 1a–b & 2a–d)**. The docked models obtained from CluPro server are further subjected to flexible refinement using FIREDOCK server(Andrusier et al., 2007; Mashiach et al., 2008). We compare the affinity of these receptors towards full spike (with missing RBM residues) and RBD alone. The Global binding energy and Atomic contact energy values are shown in **Fig 3**. While ACE2 shows higher affinity than DC-SIGN towards the spike protein, the affinity of L-SIGN is similar to ACE2 towards the nCOVID-19 spike protein. ACE2 also shows similar affinity towards both full spike and RBD alone are similar, suggesting that in the absence of RBM residues in spike, ACE2 has a good chance of binding with the NTD region. Interestingly, DC/L-SIGN show very low affinity towards RBD alone. This further strengthen the idea that residues distinct from RBD region are involved in DC/L-SIGN-Spike protein interactions. Due to distinct binding residues in NTD and RBD regions of the spike protein binding residues, DC/L-SIGN has the potential to mediate ACE2 independent nCOVID-19 entry into the cells and inhibiting the RBM region alone may not prevent viral entry.

**Fig 2:**
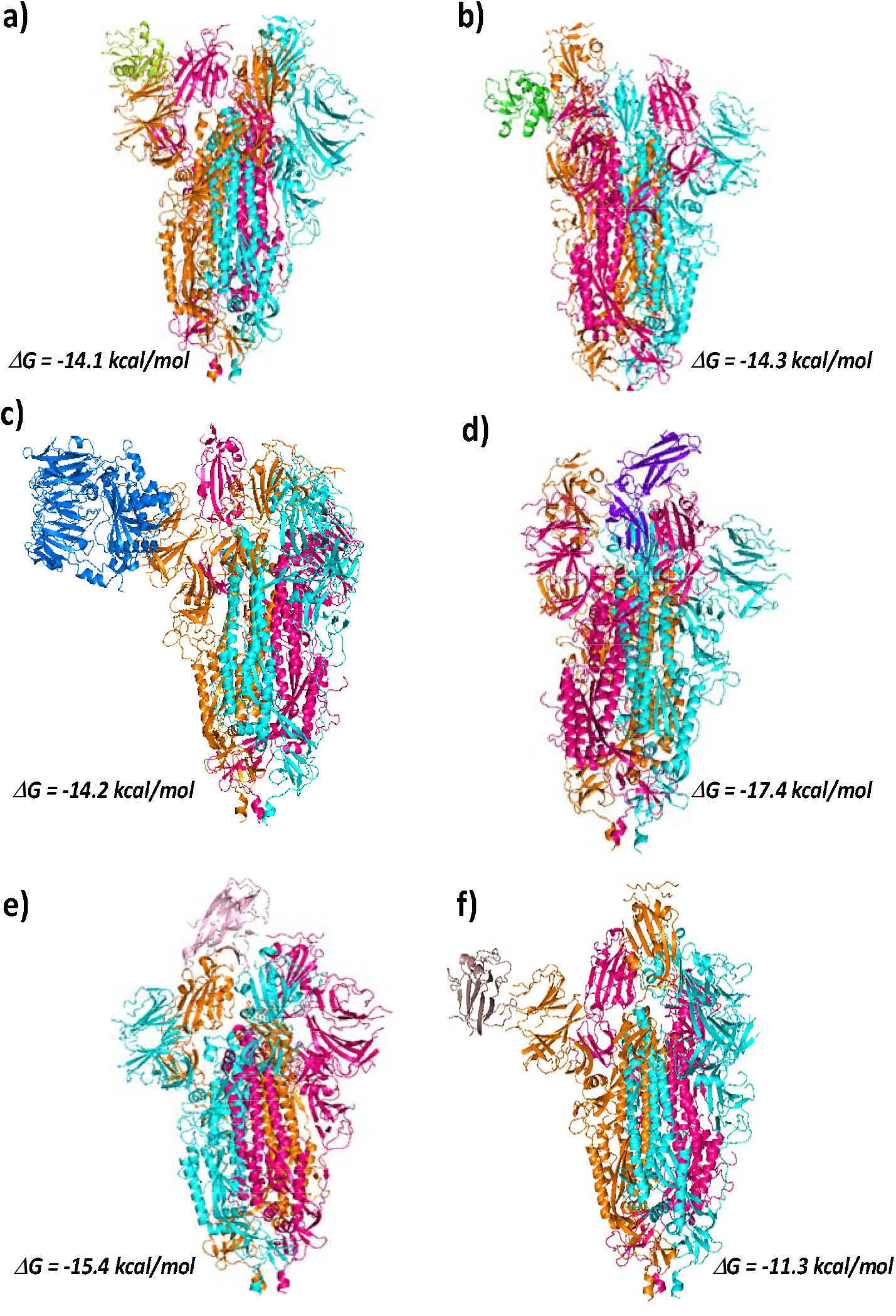
Interaction of nCOVID-19 spike protein with different cellular receptors. ClusPro models showing interaction of COVID-19 spike protein with a) DC-SIGN receptor (CRD region, in Limon) (PDB ID: 6GHV), c) L-SIGN receptor (CRD region, in Green) (PDB ID: 1XAR), c) DPP4 receptor (PDB ID: 4L72, in Blue), d) IL22 receptor (PDB ID: 3DGC, in purple), e) EphA2 receptor (PDB ID: 3C8X, in Pink)and f) EphB2 receptor (PDB ID: 3ETP, in Brown); Free energy of binding is calculated using PRODIGY server.

**Table 1a:** Hydrogen Bonds between RBD and ACE2

| Table 1a: Hydrogen Bonds between RBD and ACE2 |  |  |  |  |
| --- | --- | --- | --- | --- |
| RBD |  | ACE2 |  | Distance Å |
| ASN | 487 | GLN | 24 | 2.69 |
| LYS | 417 | ASP | 30 | 2.9 |
| TYR | 449 | ASP | 38 | 2.7 |
| THR | 500 | TYR | 41 | 2.71 |
| THR | 500 | TYR | 41 | 2.71 |
| TYR | 449 | GLN | 42 | 2.79 |
| ASN | 487 | TYR | 83 | 2.79 |
| GLY | 502 | LYS | 353 | 2.78 |

**Table 1b:** Salt Bridges between RBD and ACE2

| Table 1b: Salt Bridges between RBD and ACE2 |  |  |  |  |
| --- | --- | --- | --- | --- |
| RBD |  | ACE2 |  | Distance Å |
| LYS | 417 | ASP | 30 | 2.9 |

**Table 2a:** Hydrogen Bonds between RBD and DC-SIGN

| Table 2a: Hydrogen Bonds between RBD and DC-SIGN |  |  |  |  |
| --- | --- | --- | --- | --- |
| RBD |  | DC-SIGN |  | Distance Å |
| ASP | 389 | HIS | 254 | 2.86 |
| CYS | 361 | TRP | 258 | 2.88 |
| ASN | 360 | TRP | 258 | 2.81 |
| TYR | 365 | TRP | 258 | 2.87 |
| PHE | 392 | GLU | 259 | 2.9 |
| ARG | 357 | GLU | 259 | 2.87 |
| SER | 359 | GLU | 259 | 2.92 |
| VAL | 395 | GLU | 259 | 2.91 |
| THR | 393 | TRP | 260 | 2.75 |
| ASN | 360 | TRP | 260 | 2.88 |

**Table 2b:** Salt Bridges between RBD and DC-SIGN

| Table 2b: Salt Bridges between RBD and DC-SIGN |  |  |  |  |
| --- | --- | --- | --- | --- |
| RBD |  | DC-SIGN |  | Distance Å |
| ARG | 357 | GLU | 259 | 2.76 |

**Table 2c:** Hydrogen Bonds between RBD and L-SIGN

| Table 2c: Hydrogen Bonds between RBD and L-SIGN |  |  |  |  |
| --- | --- | --- | --- | --- |
| RBD |  | L-SIGN |  | Distance Å |
| THR | 393 | GLU | 262 | 2.82 |
| ASP | 428 | ARG | 263 | 2.76 |
| LEU | 390 | ARG | 266 | 2.64 |
| LEU | 390 | ARG | 266 | 2.7 |
| PHE | 392 | GLN | 276 | 2.89 |
| ASN | 360 | GLN | 276 | 2.68 |

**Table 2d:** Salt Bridges between RBD and L-SIGN

| Table 2d: Salt Bridges between RBD and L-SIGN |  |  |  |  |
| --- | --- | --- | --- | --- |
| RBD |  | L-SIGN |  | Distance Å |
| ASP | 428 | ARG | 263 | 2.71 |

**Fig 3:**
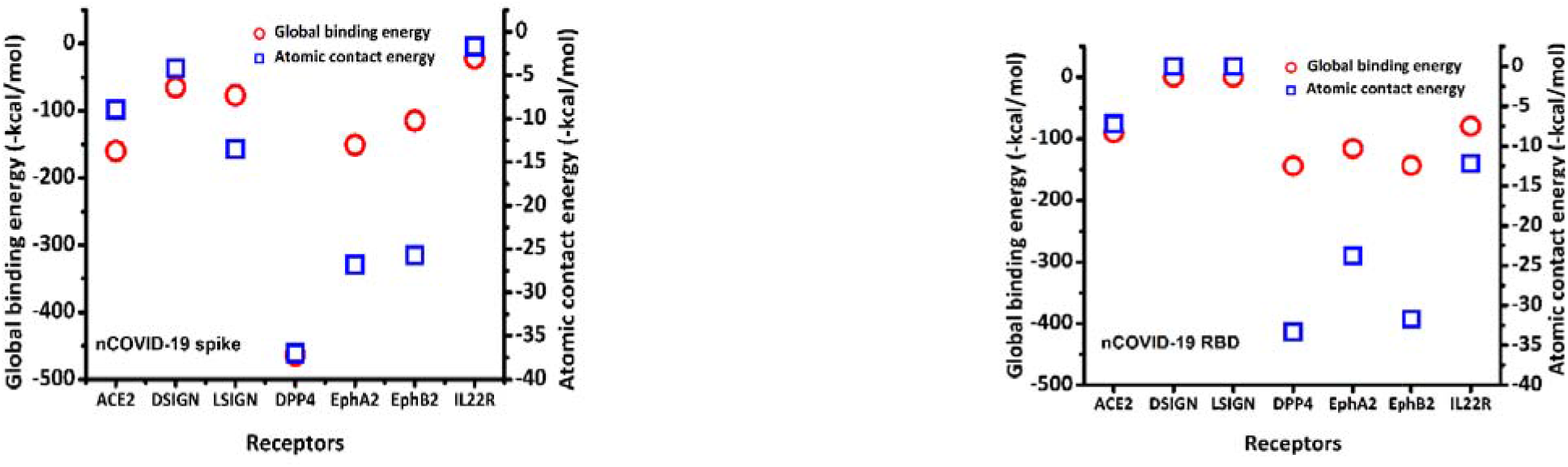
Global binding energy and Atomic contact energy for the receptor-spike and receptor-RBD docked models calculated post flexible refinement using FIREDOCK server.

### Unravelling the link between patient comorbidity and nCOVID-19 infection

A link between comorbidities and nCOVID-19 infection is evident from recent studies. One report from Italy suggests that more than 2/3^rd^ of the patients who died of COVID-19 had diabetes, cardiovascular diseases and cancer (Remuzzi & Remuzzi, 2020). Two independent studies from China indicate diabetes to be the second most common comorbidity after hypertension in patients infected with nCOVID-19 (Liang et al., 2020; Zhou et al., 2020). Alteration in the expression levels of certain cell receptors is a hallmark of diseases such as diabetes and cancer. For example, L-SIGN is reportedly overexpressed in cervical cancer and is known to promote colon cancer (Na et al., 2017; X. Wang et al., 2017). While its expression is lowered in lung cancer patients, it is found to be overexpressed in patients with brain metastasis (Liu et al., 2015). DC-SIGN is also associated with mediating the progression of gastric cancer but has a lower level of expression in patients suffering from non-Hodgkin lymphoma (Z. Zhang et al., 2013). On comparing the affinities of these receptors towards nCOVID-19 and SARS-COV spike, we observe that DC-SIGN shows higher affinity towards SARS-COV spike, while L-SIGN displays similar affinities towards both (**Table 3**). It’s therefore not clear what makes nCOVID-19 more virulent and potent danger to patients with comorbidities.

**Table 3:**
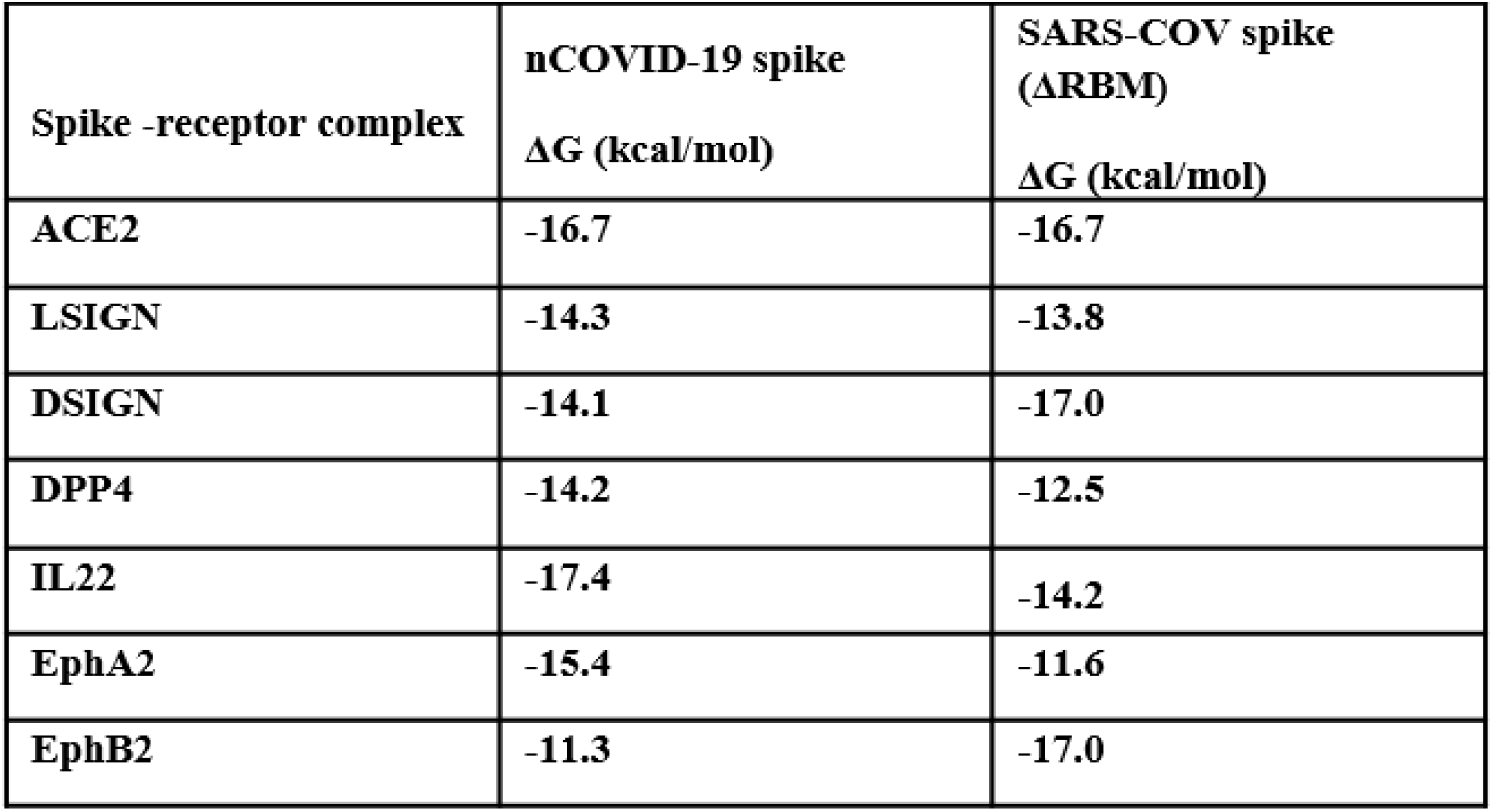
Free energy of interaction between nCOVID-19 and SARS-COV(ΔRBM) spike protein with cell receptors.

We therefore hypothesize that beside ACE2 and DC/L-SIGN receptors, certain cell receptors overexpressed in patients suffering from diabetes and cancer could also act as receptors for nCOVID-19. In the literature, we find that cell receptors such as DPP4 (Dipeptidyl peptidase-4), IL-22R (Interleukin 22 receptor) and ephrin family of receptors are overexpressed in various cancers. For example, DPP4 has been found to be overexpressed in colorectal and lung cancers (Bishnoi et al., 2019; Jang et al., 2016). DDP4 receptor also plays role in regulating insulin metabolism and has been associated with diabetes (Röhrborn, Wronkowitz, & Eckel, 2015). Interestingly, DPP4 was identified as a receptor for Human Coronavirus-Erasmus Medical Centre (hCOV-EMC) and Middle East respiratory syndrome coronavirus (MERS-CoV) (Y. Han & Král, 2020; Li et al., 2020; Raj et al., 2013). Ephrin receptors such as EphA2 and EphB2 also promote viral entry by interacting with the viral surface glycoprotein (Lee, 2007; J. Wang, Zheng, Peng, Zhang, & Qin, 2020; H. Zhang et al., 2018). IL-22R is found be overexpressed in non-small cell lung cancer (Guillon et al., 2016). However, there are no reports on the role of IL-22R receptor in viral entry into the cells.

In this section we look for the binding affinity of nCOVID-19 spike protein trimer towards extracellular domains of DPP4 (PDB ID: 4L72, chain A), IL-22R (PDB ID: 3DGC, chain R), EphA2 (PDB ID: 3C8X) and EphB2 receptors **(Fig 2c-f))**. Highest affinity is seen in case of IL-22R with a ΔG value of −17.4 kcal/mol. This is very close to the affinity seen in case nCOVID19 spike-ACE interaction. Amino acids of all the three chains of spike protein trimer are involved in H-bonding and salt bridge formation with the receptor IL-22R. The participation of all the three spike chains in binding with the receptor could be the reason for such a strong affinity. **(Supporting file, Table S6a-f)**. Receptor DPP4 binds to the chain B of spike protein (ΔG = −14.2 kcal/mol) using 11 H-bonds and 4 salt bridges (**Supporting file, Table S7a-b)**. The ephrin receptors EphA2 and EphB2 bind to the chain A and chain C of the spike protein respectively (**Supporting file, Table S8a-d)**. Lowest affinity towards the spike protein is seen in case of EphB2 (ΔG = −11.3 kcal/mol). In case of IL-22R the amino acid residues of RBD region make polar contacts with the receptors. Whereas, in case of DPP4 and ephrin receptors residues in the NTD of the spike protein make polar contact with the receptors. (**Supporting file, Table S6 & S7)**.

Due to the absence of RBM residues in the nCOVID-19 spike structure, we look for interactions of these receptors with RBD region of nCOVID-19 spike protein **(Supporting file, Fig S1c-f)**. DPP4 shows maximum affinity towards the nCOVID-19 RBD, with a ΔG value of −12.8 kcal/mol. High binding affinity of DPP4 towards the nCOVID-19 Spike RBD may enhance viral cell entry making diabetic patients more vulnerable. It is also noteworthy that in RBD-receptor docking, only ACE2 receptor interacts with the RBM region of the spike protein, while other cell receptors used in the study are predicted to interact with residues beyond the RBM region **(Table 4, 5 & 6)** This strengthen the idea that in a scenario where cell receptors other than ACE2 are involved, blocking only the RBM region with an inhibitor may not be sufficient to impede the viral cell entry. For a better comparison of the binding affinities, we perform flexible refinement of the docked models using FIREDOCK server **(Fig 3)**. Notably, receptor DPP4 shows maximum affinity towards both full spike (with missing RBM residues) and RBD alone (**Fig 3)**. Next to DPP4, Ephrin receptors EphA2 and EphB2 show the maximum affinity towards both nCOVID-19 spike and its RBD alone. Although post refinement, we observer lower affinity of IL22R towards full spike, its affinity towards RBD alone is comparable to that of ACE2. Together, these results strongly support our hypothesis and indicates the role of cell receptors linked to diabetes and cancer, in mediating nCOVID-19 cell entry. We further compare the binding affinity of SARS-COV spike protein towards DPP4, IL22R and ephrin receptors with that of nCOVID-19 spike protein. The receptors DPP4, IL22R and EphA2 show higher affinity towards nCOVID-19 spike than SARS-COV spike. However, it should be noted that except DPP4 and ACE2, other receptors show higher affinity towards SARS-COV RBD as compared to nCOVID-19 RBD (**Table 7**). This signifies that residues other than RBD in nCOVID-19 spike are crucial for spike-receptor binding.

**Table 4a:**
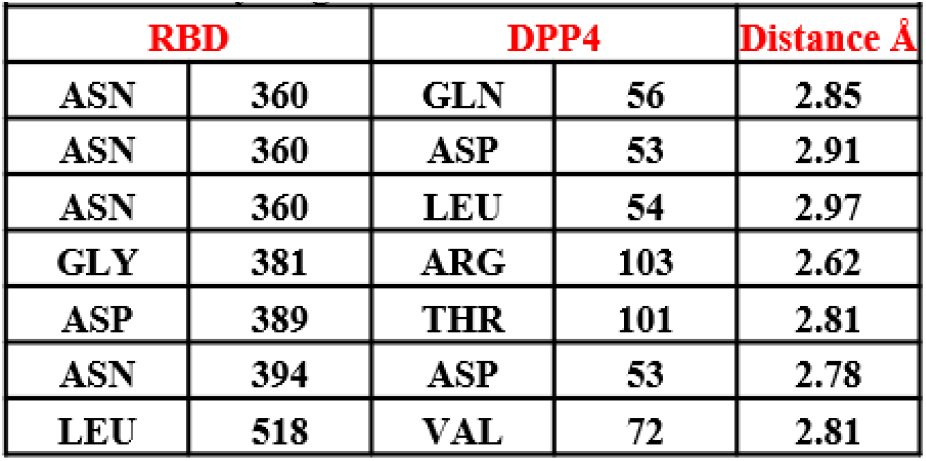
Hydrogen bonds between RBD and DPP4

**Table 4b:**
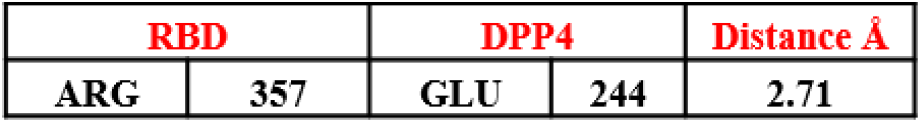
Salt Bridges between RBD and DPP4

**Table 4c:** Hydrogen bonds between RBD and IL22R

| Table 4c: Hydrogen bonds between RBD and IL22R |  |  |  |  |
| --- | --- | --- | --- | --- |
| RBD |  | IL22R |  | Distance Å |
| LYS | 386 | GLN | 117 | 2.7 |
| VAL | 362 | TRP | 208 | 2.88 |

**Table 4d:** Salt Bridges between RBD and Il22R

| Table 4d: Salt Bridges between RBD and IL22R |  |  |  |  |
| --- | --- | --- | --- | --- |
| RBD |  | IL22R |  | Distance Å |
| ASP | 389 | ARG | 112 | 2.76 |

**Table 5:** Hydrogen Bonds between RBD and EphA2

| Table 5: Hydrogen Bonds between RBD and EphA2 |  |  |  |  |
| --- | --- | --- | --- | --- |
| RBD |  | EphA2 |  | Distance Å |
| ASN | 360 | GLN | 56 | 2.85 |
| ASN | 360 | ASP | 53 | 2.91 |
| ASN | 360 | LEU | 54 | 2.97 |
| GLY | 381 | ARG | 103 | 2.62 |
| ASP | 389 | THR | 101 | 2.81 |
| ASN | 394 | ASP | 53 | 2.78 |
| LEU | 518 | VAL | 72 | 2.81 |

**Table 6:**
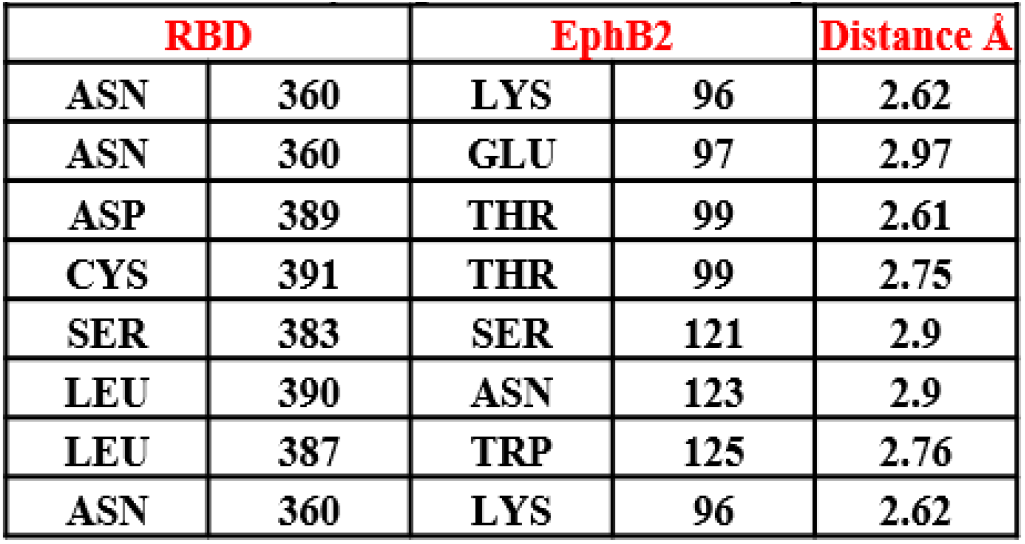
Hydrogen Bonds between EphB2

**Table 7:** Free energy of interaction between nCOVID-19 and SARS-COV RBD with cell receptors.

|  | **nCOVID-19 RBD** | **SARS-COV RBD** |
| --- | --- | --- |
| **RBD -receptor complex** | **$\Delta G$ (kcal/mol)** | **$\Delta G$ (kcal/mol)** |
| **ACE2** | **-11.9** | **-10.8** |
| **LSIGN** | **-9.8** | **-11.6** |
| **DSIGN** | **-10.7** | **-11.2** |
| **DPP4** | **-12.8** | **-11.5** |
| **IL22** | **-9.0** | **-10.9** |
| **EphA2** | **-10.8** | **-10.7** |
| **EphB2** | **-10.8** | **-11.6** |

### Structural basis for increased vulnerability

Our docking results suggests that nCOVID-19 spike is more versatile than SARS-COV spike when it comes to binding with different cell receptors. To understand the structural basis behind this versatility, we need to elucidate how flexible the nCOVID-19 spike is in comparison to SARS-COV spike. On comparing the disorderdness of both the spike protein using PONDR server (B. Xue et al., 2010), we find that 8.09% of nCOVID-19 spike and 5.84% of spike are disordered. Higher the disorderedness, higher will be the chance of a protein to interact with its binding partners. Next, we question the implications of higher flexibility in nCOVID-29 spike on its ability to bind to receptor partners. In recent years, phase separation upon protein-protein interaction has been shown to be crucial in biological functions (Feng, Chen, Wu, & Zhang, 2019). Keeping this in mind, we hypothesize that viral interactions with cellular receptors may be an event of phase separation. To understand this, we use CatGRANULE server to predict the propensity of the spike protein to undergo phase separation (Bolognesi et al., 2016). Interestingly, we find that nCOVID-19 shows higher phase separation propensity compared to SARS-COV (**Fig 4 a & b**). This suggests that nCOVID-19 spike is more fluidic in nature as compared to the SARS-COV spike. We propose that higher phase separation property of nCOVID-19 spike protein could increase its ability to interact with various cell receptors, making the virus more virulent than SARS-COV.

**Fig 4:**
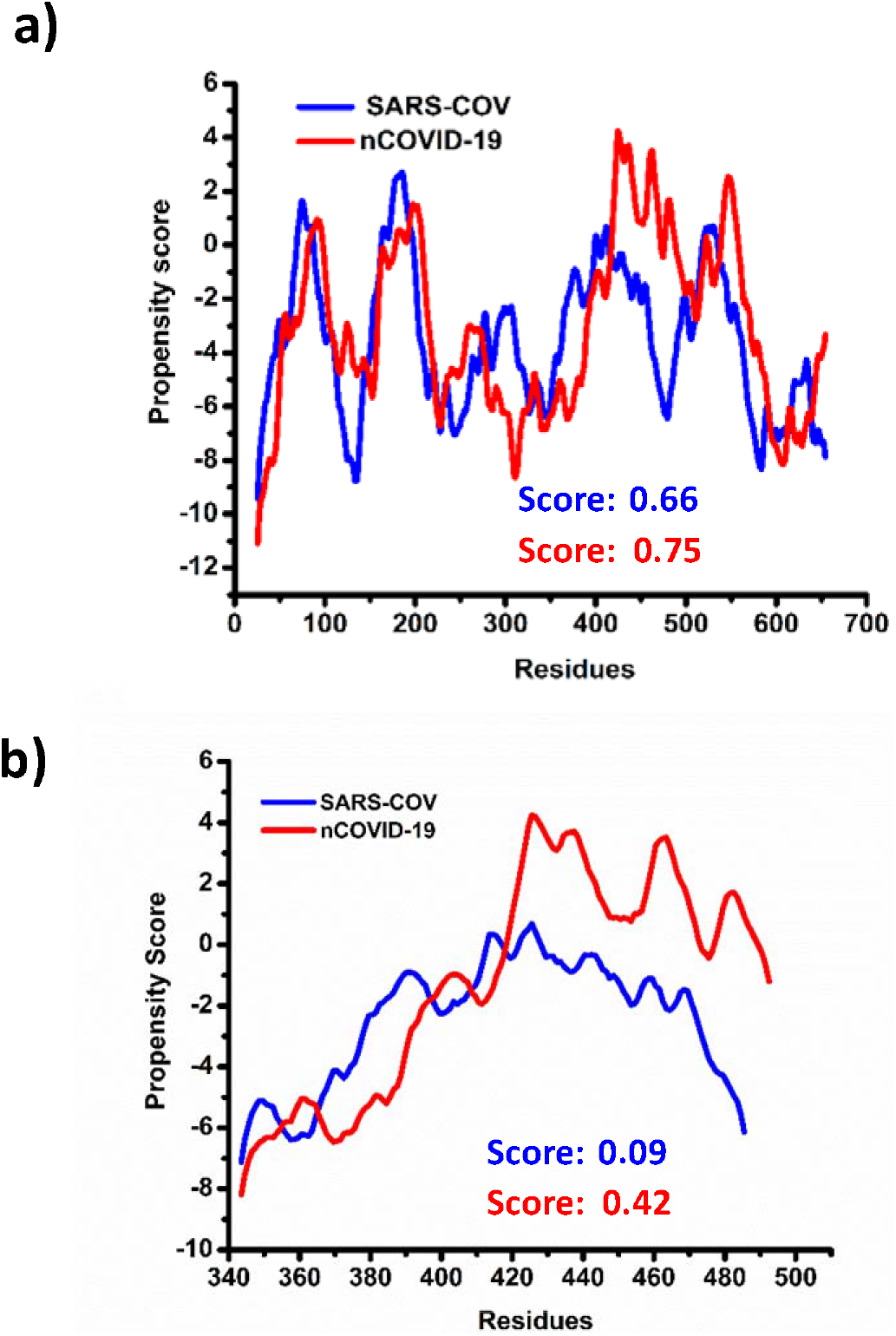
a) Phase separation propensity of S1 domain of nCOVID-19 spike protein and SARS-COV spike protein predicted using catGRANULE server. b) Phase separation propensity of nCOVID-19 RBD and SARS-COV RBD residues (340-495) predicted using catGRANULE server. The RBD region of nCOVID-19 has a higher propensity than SARS-COV RBD to undergo phase separation.

## Discussion

The ACE2 receptor is known to specifically interact with the RBD region of the spike protein and mediate viral entry into the cells. Recently, ACE2 based peptides targeting RBD region of the spike protein have been suggested to be used in peptide therapeutics against nCOVID-19. However, the affinity of ACE2 and other cellular receptors towards regions beyond the RBD of S1 domain needs to be understood. In our study we find that the association between the ACE2 and the spike protein is not impaired in the absence of the RBM residues in the spike protein. The ACE 2 receptor could bind to the NTD region of the spike protein. The implications of this interaction on the viral cell entry need experimental verification.

Earlier, receptors DC/L-SIGN have been shown to allow ACE2 independent SARS-COV entry into the cells (D. P. Han et al., 2007). The residues at positions 109, 118, 119, 158, 227 of the NTD of the spike protein were involved in mediating ACE2 independent SARS-COV entry into the cells. In the same study, the ACE2 based peptide against the spike protein could not prevent viral entry, confirming distinct binding sites of these receptors on the spike protein. In our study, we too look for the affinity of nCOVID-19 spike protein towards DC/L-SIGN receptors. Careful analysis of the interacting residues helps us to identify spike residues beyond the RBD region involved in salt bridge formation with these receptors.

There is growing evidence that links severity of nCOVID infection with patient’s comorbidities. Patients with comorbidities like diabetes and cancer are more vulnerable to nCOVID infection. We intend to understand the underlying cause behind this link by studying the affinity of nCOVID-19 spike protein with diabetes and cancer associated cell receptors. . Our docking study shows a strong affinity of nCOVID spike protein towards DPP4 receptor. DPP4 is a target for the treatment of type 2 diabetes. Strong affinity between DPP4 and RBD could explain the susceptibility of patients suffering from diabetes. Recently, Iacobellis 2020 has suggested that DPP4 inhibitors might prove to be potential players in COVID-19 therapeutics (Iacobellis, 2020). However, it must be noted that in case of hCOV-EMC, DPP4 inhibitors could not prevent DPP4 mediated viral entry into the cells (Raj et al., 2013). Beside development of new DPP4 inhibitors, synthesis of DPP4 based peptide against spike protein could be a useful strategy for nCOVID-19 treatment, However, experimental works will be required to establish a direct link between DPP4 and nCOVID-19.

Receptors such as IL-22R which are overexpressed in lung cancer must also be taken into consideration while looking for receptors targeting nCOVID-19. Although there is no evidence that could support association between IL-22R and coronavirus infection, strong affinity between the two calls for an experimental verification. We have also considered Ephrin family receptor due to their aberrant expression in various types of human cancers such as breast cancer, lung cancer, ovarian cancer, colon cancer and gastric cancer (Kikuchi et al., 2015; Kou & Kandpal, 2018). The receptor EphA2 is already reported to facilitate cellular entry of viruses such as hepatitis C virus, Kaposi’s sarcoma-associated herpesvirus (J. Wang et al., 2020; H. Zhang et al., 2018). In our study we find strong association between EphA2 receptor and nCOVID-19 spike. Further experimental work in identification of nCOVID-19 receptors may pave the way towards the development of targeted therapeutics against the virus.

## Acknowledgements

AG and GK thank INST, Mohali for fellowship.

## Authors contribution

SS, AG and GK conceived the idea. AG and GK performed the studies and analyzed the data. GK, AG, SS wrote the manuscript.

## Funding

Institutional research grant.

## Conflict of Interest

The authors declare no conflict of interest.

## Notes

### Competing Interest Statement

The authors have declared no competing interest.

