## Supporting figures and tables for "New insights into nCOVID-19 binding domain and its cellular receptors"

**Figure S1**

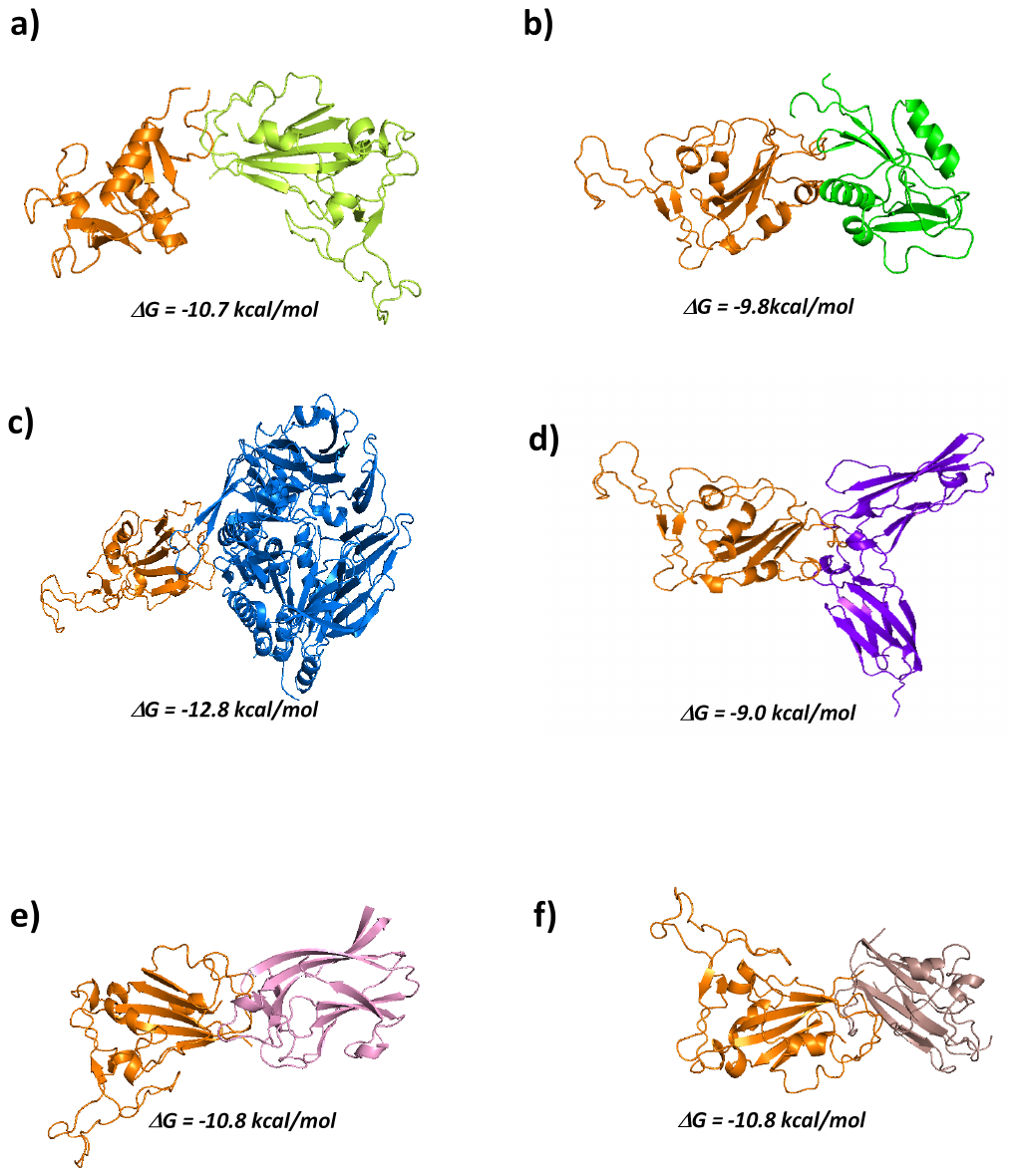

**Fig S1:** Interaction of nCOVID-19 RBD with different cellular receptors. ClusPro models showing interaction of RBD region of spike protein with a) DC-SIGN receptor (CRD region), c) L-SIGN receptor (CRD region), c) DPP4 receptor, d) IL22 receptor, e) EphA2 receptor and f) EphB2 receptor; Free energy of binding is calculated using PRODIGY server.

**Table S1**

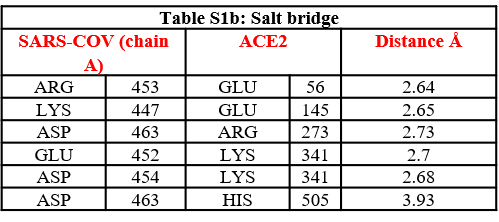

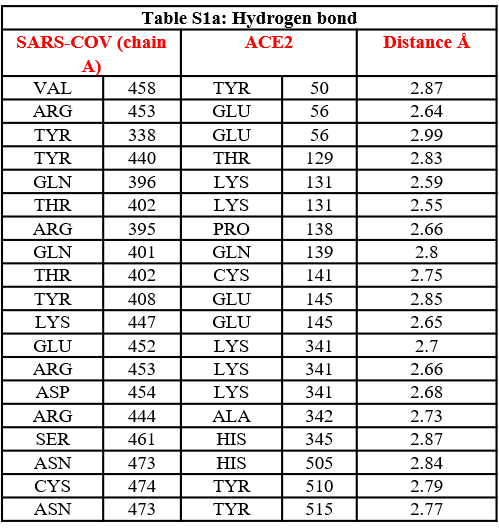

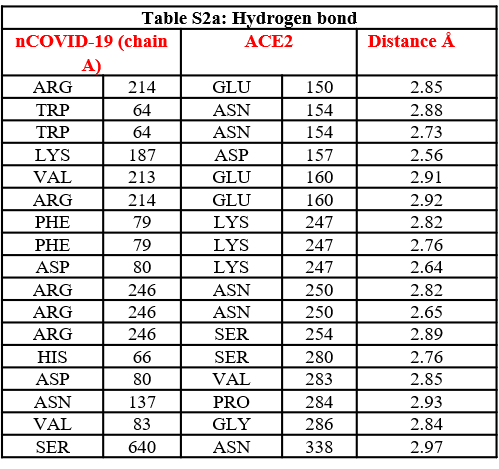
**Table S2**

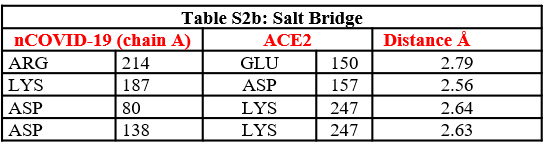

**Table S3**

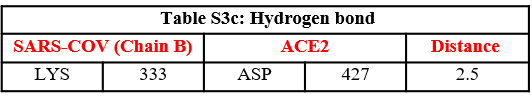

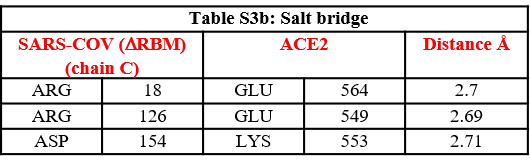

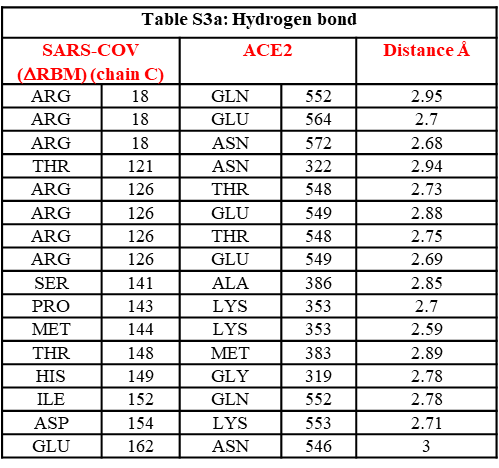

**Table S4**

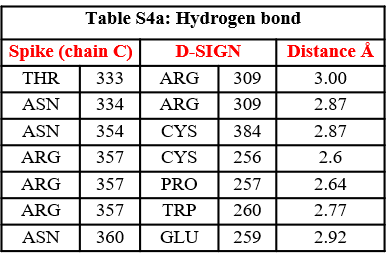

**Table S5**

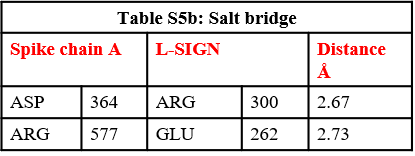

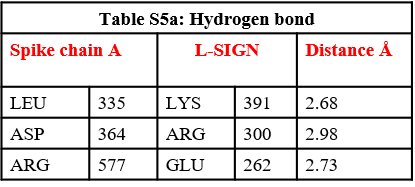

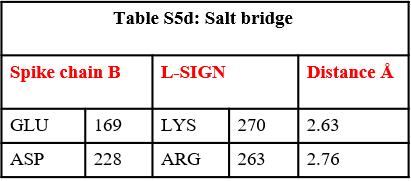

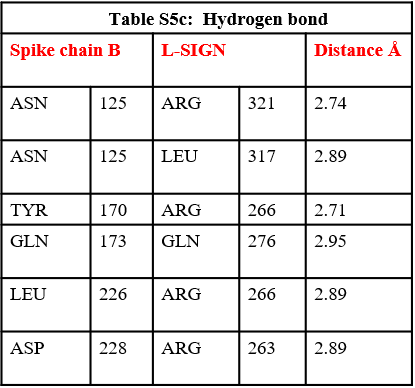

**Table S6**

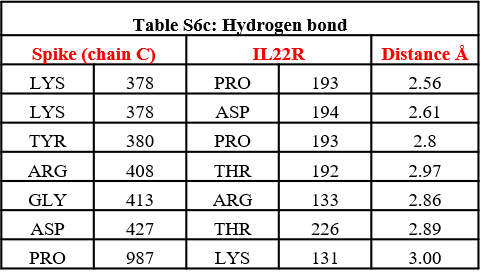

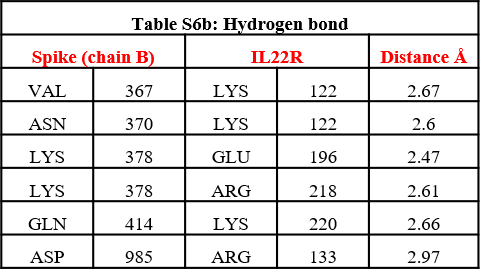

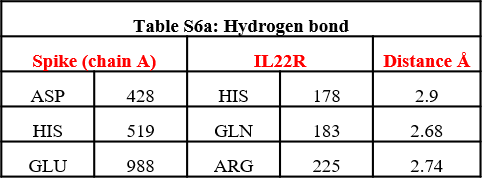

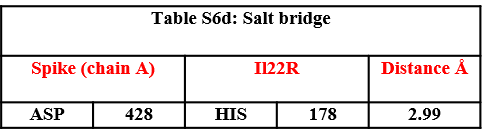

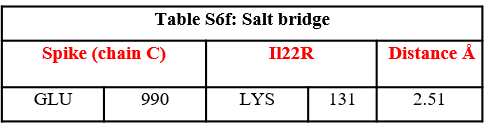

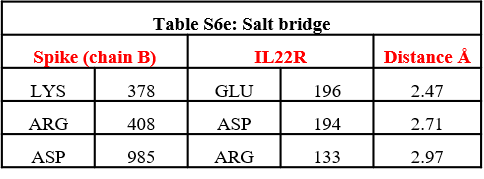

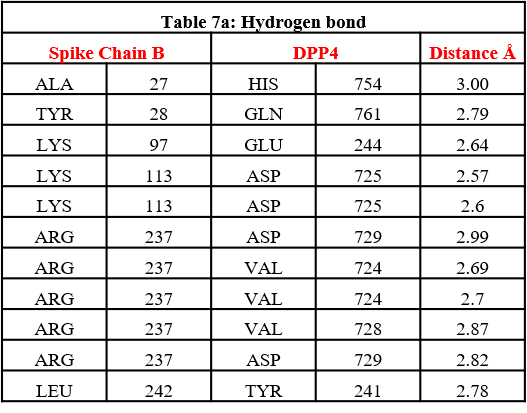
**Table S7**

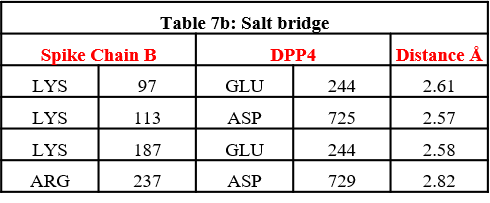

**Table S8**

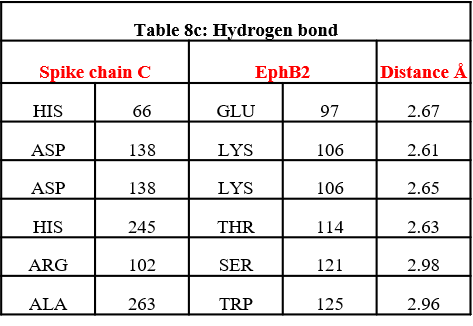

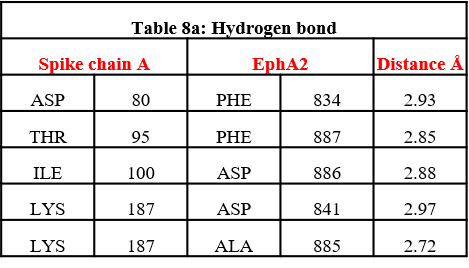

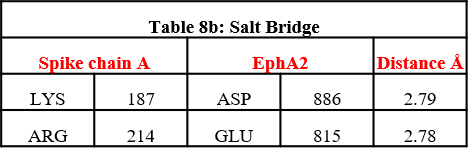

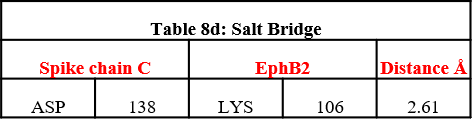
